# Dynamic interactions between epithelial skin cells and a sensory cavity sculpt the growing olfactory orifice

**DOI:** 10.1101/2025.07.25.666761

**Authors:** Gordillo Pi Clara, Cabrera Mélody, Gilles Jean-François, Bardet Pierre-Luc, Eschstruth Alexis, Bonnet Isabelle, Breau Marie Anne, Baraban Marion

## Abstract

During morphogenesis and in pathological conditions, gaps can form in the plane of epithelial barriers upon cellular forces that disrupt intercellular junctions. How the size of these epithelial holes further increases over time and what sets their shape remain poorly understood. Here we analyze the formation of the olfactory orifice (the nostril) in zebrafish, which opens and grows in the skin epithelium above a rosette of olfactory placode cells, allowing the sensory neurons to directly access odor cues. Using quantitative imaging and tissue-specific perturbations, we analyzed the dynamic remodeling of skin cells allowing the expansion of the orifice edge. We identified the sensory cavity located in the center of the placodal rosette as a crucial player that sets the size of the growing epithelial hole in the skin. We further showed that fine-tuning of actomyosin contractility within each tissue (skin and sensory cavity) exerts non-autonomous effects on the neighboring tissue, thereby shaping the nostril structure. This study uncovers dynamic cell behaviors and reciprocal tissue-tissue interplay that control the growth and shape of an epithelial hole *in vivo*.

## Introduction

During embryonic development and adult life, epithelial tissues may undergo extensive morphological rearrangements involving a partial loss of epithelial integrity. One example of such epithelial barrier remodeling is the formation of gaps in the plane of epithelia^1^, referred to as planar epithelial holes. These holes can be distinguished from lumen formation, where enclosed, fluid-filled cavities open on the apical side of epithelial sheets, and from epithelial folding initiating the morphogenesis of tubular structures^2,3^. Planar epithelial holes have been reported to form *in vitro* and *in vivo*, and display diverse functions in morphogenesis, adult physiology and pathological conditions^1^. For instance, a transient opening of multiple tricellular junctions in the follicular epithelium is crucial for the maturation of the Drosophila egg^4–6^. At a larger scale, the rapid and elastic deformation of the mouth allows Hydra to feed^7^, and holes and fractures observed upon body movements in *Trichoplax adhaerens* trigger morphological changes that facilitate asexual reproduction^8^. Permanent ruptures of epithelial layers are also observed *in vivo* during follicular and peripodial epithelia breakdowns, which are required for ovulation and leg growth, respectively^9–11^, and *in vitro* when extrinsic forces are applied^12–15^. Round and smooth holes can also spontaneously form upon epithelial contractions and/or low adhesion to the substrate in culture^9,16^. The mechanisms driving the opening of planar epithelial holes are starting to be well understood. Because they form in the plane of epithelial sheets, their opening does not rely on osmotic pressure as seen in luminogenesis, but rather on cellular mechanical forces allowing the tearing and unzipping of intercellular junctions, which build up within the epithelium itself^7–11,16^, or are extrinsic to the epithelium^17,18^. However, how planar holes continue to expand in the epithelial layer and how their shape is established, remain to be elucidated.

Here we analyze the growth of an epithelial hole, the olfactory orifice, which opens in the skin epithelium above the olfactory placode in zebrafish. This hole prefigures the future nostril and is crucial for the olfactory function as it exposes the sensory neurons to external odors. Before its opening, a subset of olfactory placode cells assembles into a rosette structure beneath the skin cells. Their apical tips line a semi-ellipsoidal cavity in the rosette center, which is in contact with the skin epithelium^17^. We previously found that the olfactory orifice opens in the skin upon pulling forces exerted by the rosette neurons^17^, thus exposing the sensory neurons through this cavity (Fig. 1a). In the present study we combined live imaging, laser ablation, molecular tools and tissue-specific perturbations to dissect the mechanisms regulating the coordinated growth and shaping of the epithelial hole and the underlying placodal cavity. We analyzed the cell behaviors involved in the enlargement of the two structures and identified placodal cavity growth as a main driving force for orifice expansion in the skin. Furthermore, a multicellular ring of actomyosin located in the upper part of the cavity acts non-autonomously to smoothen the edge of the growing orifice in the skin. In turn, epithelial skin cell relaxation downstream of ERK signaling is important to maintain the open shape of the placodal cavity. This work identifies key cell behaviors, reciprocal tissue-tissue interactions and molecular players driving the growth and shaping of a planar epithelial orifice *in vivo*.

**Figure 1.**
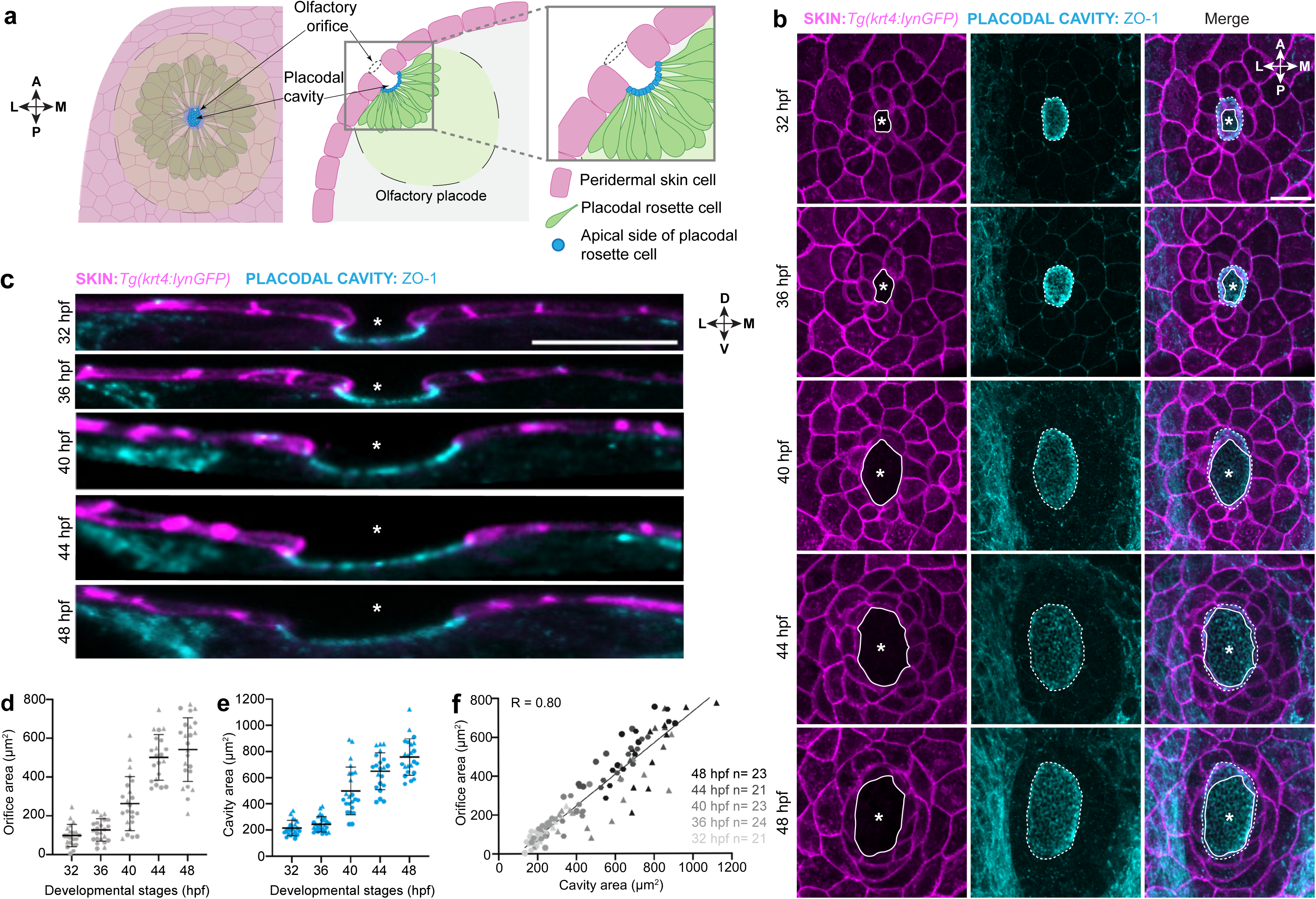
Coordination between orifice and cavity growth. **a.** Schematic view of the olfactory orifice in the skin and the rosette of olfactory placode cells right after the opening of the orifice at 32 hpf. Dorsal view on the left, and cross section on the right. **b.** Time course analysis. *Tg(krt4:lynGFP)* embryos, in which the membrane of peridermal skin cells are labelled, were fixed every 4 hours and immunostained for GFP (magenta) and for the tight junction protein ZO-1 (cyan) to reveal the apical tips lining the cavity. Dorsal views, maximum projections. Asterisk = orifice. White line = orifice contour. Dotted line = cavity contour. Scale bar = 20 µm. **c.** Orthogonal views of the embryos shown in b. Asterisk = orifice. Scale bar = 20 µm. **d, e**. Quantification of orifice (d) and cavity (e) areas (3 independent experiments; 32 hpf: n = 21 embryos; 36 hpf: n = 24 embryos; 40 hpf: n = 23 embryos; 44 hpf: n = 21 embryos; 48 hpf: n = 23 embryos). **f.** Correlation between the areas of the orifice and rosette cavity, the data were pooled from all stages. Simple linear regression.

## Results

### The olfactory orifice and the placodal cavity exhibit coordinated growth

The olfactory orifice opens in the skin above the olfactory placode cavity between 28 and 34 hours post fertilization (hpf)^17^. To understand how the hole further grows in the epithelium, we used a time course analysis to characterize the morphological changes of the orifice and the underlying placodal cavity during the first 16 hours (h) of expansion (from 32 to 48 hpf, referred to as the initial expansion phase, see Supplementary Movie 1 for 3D visualization at 48 hpf). Embryos expressing the *Tg(krt4:lynGFP)* transgene to visualize the membrane of peridermal skin cells^19^ were fixed every 4 hours and immunostained for the tight-junction protein ZO-1 to reveal the apical tips of the cells lining the cavity (Fig. 1b, c). We observed a progressive growth of the orifice and cavity, both in the anteroposterior and mediolateral dimensions (Fig. 1b-e). The areas of the cavity and orifice exhibited a strong correlation (Fig. 1f), showing that they grow concomitantly, and suggesting the existence of a scaling mechanism between the two structures. To further understand how the epithelial hole grows, we next examined the dynamic behaviors of skin cells located on the edge of the orifice.

### A dynamic remodeling of skin cells takes place on the edge of the growing orifice

To monitor skin cell behaviors, we performed quantitative live imaging on *Tg(krt4:lynGFP)* embryos from 32 hpf during 14 hours (Supplementary Movie 2 and Fig. 2a, a’). A progressive increase of the orifice area and perimeter was observed (Fig. 2b, b’), consistent with our observations on fixed embryos. We noticed that the number of skin cells located close to the orifice increases over time due to mitotic events (Fig. 2a, a’, b’’, c). To analyze the spatiotemporal distribution of cell divisions, we compiled maps of all division events occurring in the field of view during the movies (Fig. 2c’ and Supplementary Fig. 1a). Divisions were enriched in the first row (bordering cells) and second row of cells neighboring the orifice compared to the rest of the epithelium (Fig. 2c’, c’’), and occurred mainly tangentially to the edge of the growing orifice (Fig. 2c’, c’’’).

**Figure 2.**
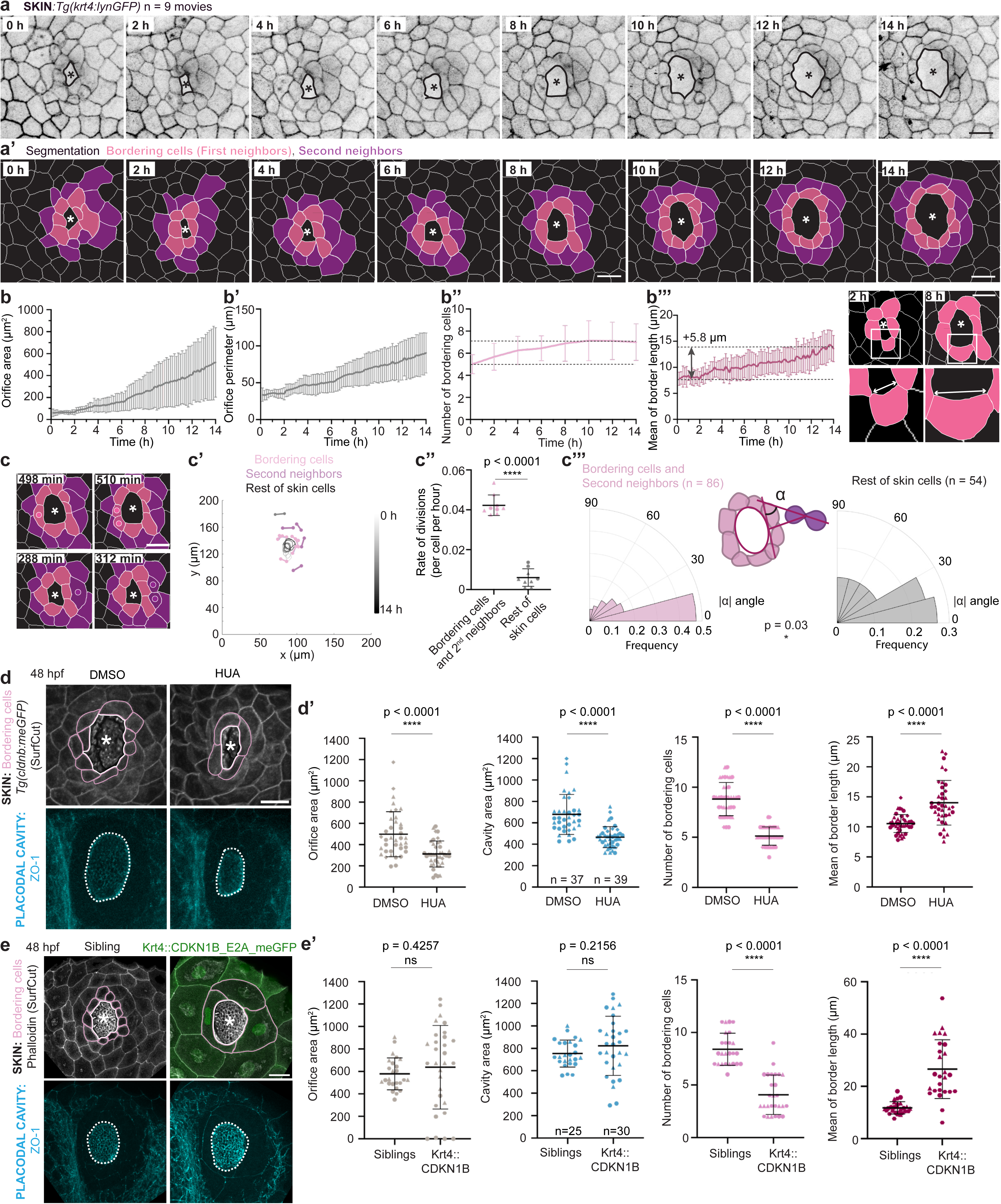
Remodeling of the skin cells around the orifice. **a.** Live imaging on a *Tg(krt4:lynGFP)* embryo, in which the membrane of peridermal skin cells are labelled (black). Dorsal view, anterior on the top, lateral on the left, maximum projections. Asterisk = orifice. Black line = orifice contour. Scale bar = 20 µm. **a’**. Segmentation of the skin cell contours. Asterisk = orifice. Pink = first neighbor skin cells (i.e. cells located on the orifice edge, or bordering cells). Purple = second neighbors. **b, b’, b’’, b’’’**. Quantification of orifice area (b), orifice perimeter (b’), number of bordering cells (b’’), and average border length (b’’’), defined as the length of the portion of bordering skin cell membranes contributing to the orifice edge, as depicted with double headed arrows in the images (right panel) extracted from the segmented movie shown in a’ (n = 9 movies from 2 independent experiments). **c.** Examples of cell divisions occurring in the bordering skin cells (pink) or second neighbors (purple), images extracted from the segmented movie shown in a’. Asterisk = orifice. The mother cell and the two daughter cells are indicated with colored circles. **c’**. Map of cell divisions occurring in the field of view during the time lapse sequence (one embryo). Grey lines = contours of the growing orifice, the time is color coded. For each division, a segment links the centers of the two daughter cells, which are represented by the dots. Pink segments = bordering skin cell divisions, purple segments = 2nd neighbor cell divisions, grey segments = divisions in the rest of the skin epithelium (for those, time is color coded). **c’’**. Quantification of the rate of cell divisions in the bordering cells and 2nd neighbors versus in the rest of the epithelium. Mann-Whitney test. **c’’’**. Rose plots showing the distribution of the absolute cell division angles (alpha) with respect to the edge of the orifice (as depicted on the schematic view, see also Methods) in the bordering cells and 2nd neighbors (pink) and the rest of the epithelium (grey). A permutation test was performed (see Methods). **d.** Images of *Tg(cldnb:meGFP)* embryos treated with DMSO or HUA from 32 hpf, fixed at 48 hpf and immunostained for GFP (grey) and ZO-1 (cyan). Top images = skin seen with the *Tg(cldnb:meGFP)* transgene and extracted with the SurfCut plugin (see Methods). Asterisk = orifice. White line = orifice contour. Bordering cells are outlined with a pink line. Bottom images = cavity seen with ZO-1 immunostaining, maximum projections. Dotted white line = cavity contour. Scale bar = 20 µm. **d’**. Quantification of orifice area, cavity area, number of bordering cells and mean border length in HUA-treated embryos and DMSO-treated controls at 48 hpf (3 independent experiments, n = 39 HUA- treated embryos, n = 37 DMSO controls). Mann-Whitney tests for orifice and cavity area, Welch’s t-tests for number of bordering cells and mean of border length. **e.** Images of *Tg(krt4:KalTA4; uas:cdkn1b-E2A-GFP)* embryos, in which cell divisions are inhibited in the skin, and control siblings fixed at 48 hpf, immunostained for ZO-1 (cyan) and stained with phalloidin-rhodamine (grey), GFP in green. Top images = skin extracted with the SurfCut plugin (see Methods). Asterisk = orifice. White line = orifice contour. Bordering cells are outlined with a pink line. Bottom images = cavity seen with ZO-1 immunostaining, maximum projections. Dotted white line = cavity contour. Scale bar = 20 µm. **e’**. Quantification of orifice area, cavity area, number of bordering cells and mean border length in *Tg(krt4:KalTA4; uas:cdkn1b-E2A-GFP)* embryos and control siblings at 48 hpf (3 independent experiments, n = 30 double transgenic embryos, n = 25 control siblings). Unpaired, two-tailed t-test for number of bordering cells, Welch’s t-tests for the other graphs.

Bordering cells also appeared to change area and shape during orifice expansion (pink cells, Fig. 2a’): they decreased their area overtime (Supplementary Fig. 1b) and by the end of the imaging period (at 46 hpf), their surface was smaller than surrounding skin cells (Supplementary Fig. 1b’), likely due to the accumulation of cell divisions in this region. At 46 hpf, bordering cells were more elongated than at 32 hpf, and along with second neighbors, they were also more elongated than surrounding skin cells, as shown by cell aspect ratio quantification (Supplementary Fig. 1b). To further quantify cell shape changes in relation with the expansion of the orifice periphery, we measured the portion of bordering skin cell membranes contributing to the orifice (referred to as border length, Fig. 2b’’’). Strikingly, the bordering cells devoted more and more length to the orifice edge (average increase of 5.8 µm +/- 0.9, Fig. 2b’’’) despite their reduction in area. Of note, recruitment of new skin cells on the orifice edge, or departure of cells from the edge occurred very rarely (2 and 9 cells, respectively, across 9 movies), ruling out neighbor exchange as an important driver of orifice growth. In conclusion, a dynamic remodeling of skin cells occurs in the vicinity of the orifice during its expansion, including tangentially-oriented cell divisions and changes in cell area and morphology.

### Skin cell divisions are not required for orifice expansion

Tangentially-oriented divisions of skin cells around the orifice could facilitate its growth by providing new cells^20^ or increasing locally the fluidity of the epithelium^21,22^. Forces exerted by mitotic rounding have also been involved in lumen expansion in the zebrafish otic placode^23^. To test the role of cell divisions in the expansion of the olfactory orifice, we pharmacologically blocked cell proliferation with hydroxyurea and aphidicolin (HUA) during the initial expansion phase, which led to a reduction of the number of bordering skin cells (Fig. 2d, d’), showing treatment efficiency. This treatment resulted in a significant decrease in the area of the olfactory orifice at 48 hpf, but the size of placodal cavity was also reduced (Fig. 2d, d’). This indicates that cell divisions in the skin and/or the olfactory placode are required for orifice expansion. Interestingly, the few remaining bordering skin cells were highly deformed, with an increased border length (Fig. 2d, d’). This suggests that cell deformation partially compensates for the reduced skin cell number in this condition, allowing a partial growth of the orifice.

To specifically test the function of cell divisions in the skin, we used skin expression of a degradation-resistant mutant of human CDKN1B (cyclin-dependent kinase inhibitor 1B), known to block divisions in MDCK cells^24^. We generated a transgenic *Tg(uas:cdkn1b-E2A- GFP)* line in which the CDKN1B mutant is placed under the control of UAS regulatory sequences. To achieve expression in the skin without expression in the placode, we produced the *Tg(krt4:KalTA4)* driver, obtained by Crispr/Cas9-mediated replacement of eGFP by KalTA4 in the *Et(krt4:EGFP)* line^25^. In *Tg(krt4:KalTA4; uas:cdkn1b-E2A-GFP)* embryos, the skin cells were larger than in control siblings (Fig. 2e), as previously shown upon inhibition of cell divisions in the periderm^26^. Unexpectedly, orifice growth occurred normally in most cases (Fig. 2e’), showing that cell divisions in the skin are dispensable for the initial expansion phase. In very few embryos (4/30), the orifice did not open, likely due to the presence of large, obstructing skin cells just above the cavity (the orifice only opens above the cavity at multiple vertices, as described in^17^). The large bordering skin cells appeared capable of drastic deformations: individual cells often contributed substantially to the orifice edge with a highly increased border length covering up to half of its perimeter (Fig. 2e, e’). Thus, in the absence of cell divisions in the skin, the orifice copes by growing through the deformation of bordering cells. Altogether, these findings show that skin cell divisions are not necessary for orifice expansion and rather suggest an important contribution of skin cell shape changes. Of note, these data also suggest that the reduced orifice size observed in HUA-treated embryos is mainly due to the reduced cavity size, which could result from a role of cell divisions in the olfactory placode. This prompted us to investigate the mechanisms of the placodal cavity growth and its contribution to the expansion of the epithelial orifice.

### The cavity grows through the intercalation of new placodal cells

The orifice in the skin and the placodal cavity grow in coordination (Fig. 1b). To assess how the cavity grows, we analyzed the apical surfaces of the placodal cells lining the cavity. At 48 hpf, by the end of the initial expansion phase, high-resolution imaging of ZO-1 immunostaining revealed two distinct categories of apical areas in the placodal cavity: small, circular tips distributed throughout the cavity and presumably belonging to the sensory neurons^17^, but also large, cuboidal apical areas specifically located in the lateral region of the cavity and likely belonging to multiciliated cells (MCCs), which are important at later stages to modulate olfaction^27^ (Fig. 3a, a’, a”, b). We investigated in detail how the two types of apical areas evolve during orifice expansion.

**Figure 3.**
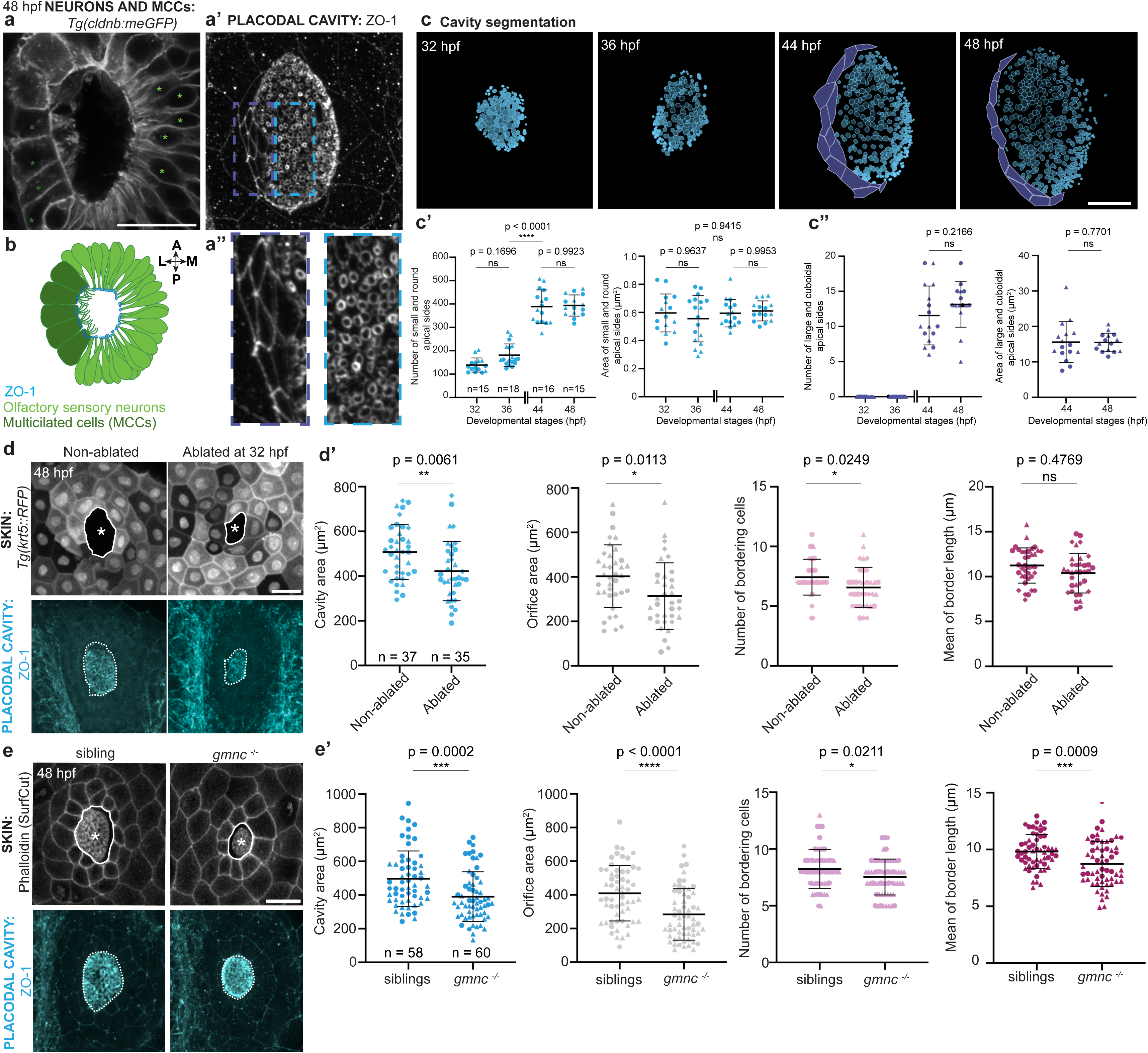
Mechanisms of cavity growth and its influence on orifice expansion. **a.** Image of a *Tg(cldnb:meGFP)* embryo at 48 hpf, showing olfactory sensory neurons (light green asterisks), and multiciliated cells (MCCs, dark green asterisks) on the lateral side of the cavity. Dorsal view, 1 z-section. Scale bar: 20 µm. **a’.** High resolution image of ZO-1 immunostaining at 48 hpf, showing the two types of apical areas contributing to the cavity: small, circular apical tips that likely belong to the sensory neurons, and large, cuboidal apical areas in the lateral region of the cavity, belonging to the MCCs. **a’’**. Zoom on the MCC apical areas (purple rectangle) and on the small and round apical areas (blue rectangle). **b.** Schematic view of the placodal rosette at 48 hpf, with olfactory sensory neurons and MCCs. **c.** Segmentation of the apical areas of the cavity, obtained from high resolution images of ZO-1 immunostaining at different stages, showing the small, round apical areas (blue) and the large cuboidal apical areas of the MCCs (purple). Scale bar = 20 µm. **c’.** Quantification of the number of small, circular apical tips and of their average area (2 independent experiments, 32 hpf: n = 15 embryos; 36 hpf: n = 18 embryos; 44 hpf: n = 16 embryos; 48 hpf: n = 15 embryos). Welch’s ANOVA followed by Dunnett T3 multicomparison test. **c’’**. Quantification of the number of large apical areas from MCCs and of their average area (2 independent experiments, 32 hpf: n = 15 embryos; 36 hpf: n = 18 embryos; 44 hpf: n = 16 embryos; 48 hpf: n = 15 embryos). Mann-Whitney tests. **d.** Images of 48 hpf placodes in which laser ablation of rosette cells was performed at 32 hpf, and of control, non ablated placodes. The *Tg(omp:meYFP)* transgene was used to label placodal cells (not shown). The *Tg(krt5:Gal4-ERT; uas:RFP)* line (grey, top) was used to visualize the orifice and ensure that the skin is not affected by the ablation (see also Supplementary Movie 4). ZO-1 immunostaining (cyan, bottom) was carried out to reveal the cavity. Scale bar = 20 µm. **d’**. Quantification of cavity area, orifice area, number of bordering cells and mean border length at 48 hpf (4 independent experiments, n = 35 ablated placodes, n = 37 control, non ablated placodes). Unpaired, two-tailed t-tests. **e.** Images of *gmnc^-/-^*mutant embryos and of control siblings at 48 hpf. Top images = skin seen with phalloidin staining (grey) and extracted with the SurfCut plugin (see Methods). Bottom images = ZO-1 immunostaining (cyan) to reveal the cavity. Scale bar = 20 µm. **e’.** Quantification of cavity area, orifice area, number of bordering cells and mean border length at 48 hpf (3 independent experiments, n = 60 *gmnc^-/-^* mutants, n = 58 control siblings). Mann-Whitney test for cavity area, unpaired, two-tailed t-tests for orifice area and number of bordering cells and Welch’s t-test for the mean border length.

Deep learning tools were used to segment the small and round apical tips and quantify their number and size at different stages of orifice expansion (Fig. 3c and Supplementary Movie 3). The number of these tips increased over time, while their mean area remained relatively constant (Fig. 3c, c’). This suggests that new cells coming from more basal regions of the placode, where progenitors have been proposed to be located at earlier stages^28^, progressively intercalate to contribute to the rosette and the cavity border. Some of these intercalation events were visible upon live imaging of sensory neuron behavior during the expansion (Supplementary Movie 4).

The large cuboidal apical areas appearing on the lateral side of the cavity are presumed to belong to MCCs^27^. To confirm their identity, we used immunostaining for Centrin, known to be highly enriched in other types of MCCs due to their multiple cilia (see for instance^29^) (Supplementary Fig. 2a). At 32 hpf, just after the opening, Centrin-positive MCCs were detected close to the cavity showing that they are already specified at this stage, however their characteristic large apical surfaces were not yet visible with the ZO-1 immunostaining (Supplementary Fig. 2a and Fig. 3c). The number of Centrin-positive MCCs increased over time during orifice expansion (Supplementary Fig. 2a’). The large apical areas started to be visible from 44 hpf onwards on the lateral side of the cavity through ZO-1 immunostaining (Fig. 3c), and their number and size did not vary between 44 and 48 hpf (Fig. 3c’’). From these findings, we conclude that the placodal cavity grows through at least two mechanisms: the addition of small apical tips through intercalation of new cells in the rosette, and the emergence of large apical tips from the MCCs, which intercalate and differentiate in the lateral cavity between the skin and the other placodal cells.

### The growth of the placodal cavity regulates the expansion of the orifice in the skin

To determine the impact of the cavity growth on the expansion of the epithelial orifice, we used two types of placode-specific perturbations. We first performed laser ablation of rosette placodal cells at 32 hpf in embryos carrying the *Tg(omp:meYFP)* transgene, which labels ciliated sensory neurons^30^ and at least a subset of MCCs^31^. The *Tg(krt5:Gal4-ERT; uas:RFP)* line was used to visualize the skin. The ablation resulted in targeted, partial destruction of rosette placodal cells (affecting potentially both neurons and MCCs) while preserving skin integrity (Fig. 3d and Supplementary Movie 5). At 48 hpf, as expected, the cavity area was significantly smaller in the ablated placodes (by around 25% in average), confirming effective placodal cell depletion. Strikingly, we observed a scaled reduction of the olfactory orifice area in the skin (Fig. 3d, d’), demonstrating the role of rosette placodal cells in setting up the size of the olfactory orifice.

We observed that large apical areas from the MCCs significantly contribute to the placodal cavity from 44 hpf onwards. To further study MCC function in orifice expansion, we examined *gmnc^sq^*^34^ mutants. Gmnc (geminin coiled-coil domain containing) is crucial for MCC differentiation, in particular to induce the multiciliation program^32,33^. At 48 hpf *gmnc* homozygous mutants displayed a smaller placodal cavity as compared to siblings, as expected (Fig. 3e, e’). This was associated with a scaled reduction in the size of the orifice in the skin (Fig. 3e, e’), reinforcing the idea that the growth of the placodal cavity is important to control orifice expansion. In both placodal perturbations (laser ablation of placodal cells and *gmnc* mutant), a slight but statistically significant decrease in the number of bordering cells was observed, as well as a reduced border length in *gmnc* mutants (Fig. 3d’, e’), suggesting that the placodal rosette influences, to a certain extent, the division and deformation of the overlying bordering skin cells.

MCCs bear multiple motile cilia that drive fluid movement in the nostril cavity, which is crucial for olfaction at later stages^27^. To analyze whether these cilia or their movement are required for orifice expansion, we used mutants in which cilia are not motile (*dnaaf1^tm^*^317b^ mutants^34,35^) or absent (*traf3ip1^tp^*^49d^*/elipsa* mutants lacking both primary and motile cilia^34,36^). In both cases, no significant difference was detected in orifice and cavity areas between homozygous mutants and control siblings (Supplementary Fig. 2b, b’, b’’, c, c’, c’’), showing that cilia are not essential for the determination of orifice size. This suggests that the presence of MCC’s apical areas, rather than their ciliary function, is necessary for the expansion of the orifice. Altogether our findings demonstrate that the growth of the placodal cavity, through the progressive addition of new apical surfaces, is a main driver of orifice expansion. Importantly, throughout the expansion phase the tight-junction-mediated contacts between the bordering skin cells and the upper part of the placodal cavity were maintained (Supplementary Movie 3), as previously shown during the opening of the orifice^17^, suggesting that the growing cavity serves as a physical scaffold for the expanding epithelial orifice.

### Actomyosin contractility in rosette neurons is required to smoothen the edge of the growing orifice in the skin

We previously showed that actomyosin-mediated pulling forces exerted on the skin by rosette neurons are necessary for the initial opening of the orifice^17^. During the opening, we reported two main actomyosin structures in the placode cavity: a multicellular actomyosin network around the cavity, the actomyosin cup, and dynamic radial cables in the placodal cells, proposed to be responsible for pulling on skin cells^17^. To explore whether actomyosin contractility is involved in orifice expansion, we analyzed non-muscle Myosin-II and F-actin localization using the *Tg(actß2:myl12.1-GFP)* reporter^37^ and phalloidin staining, respectively. No clear radial cable could be detected, except in the early phase of the expansion, just after the opening (32 hpf). However, the multicellular actomyosin cup persisted throughout orifice enlargement, with an enrichment in the apical, upper part of the cavity, forming a ring where the cavity and the bordering skin cells are in close contact (Fig. 4a and Supplementary Movie 6). From 40 hpf onward, this actomyosin ring appeared to localize deeper along the lateral side of the cavity, likely due to the presence of MCCs, which expand their apical areas between the skin and the other placodal cells (Supplementary Movie 6). In addition, similar to what was observed during the opening of the orifice, actomyosin was present at the cortex of skin cells and in microridges^38,39^ (Supplementary Movie 6). To assess whether the actomyosin ring located in the apical region of the cavity is under tension, we performed laser ablation of the ring in *Tg(actß2:myl12.1-GFP)* embryos^37^ during orifice expansion. We observed an immediate recoil following the ablation in most cases (Fig. 4b, Supplementary Movie 7), indicating that the ring is under tension.

**Figure 4.**
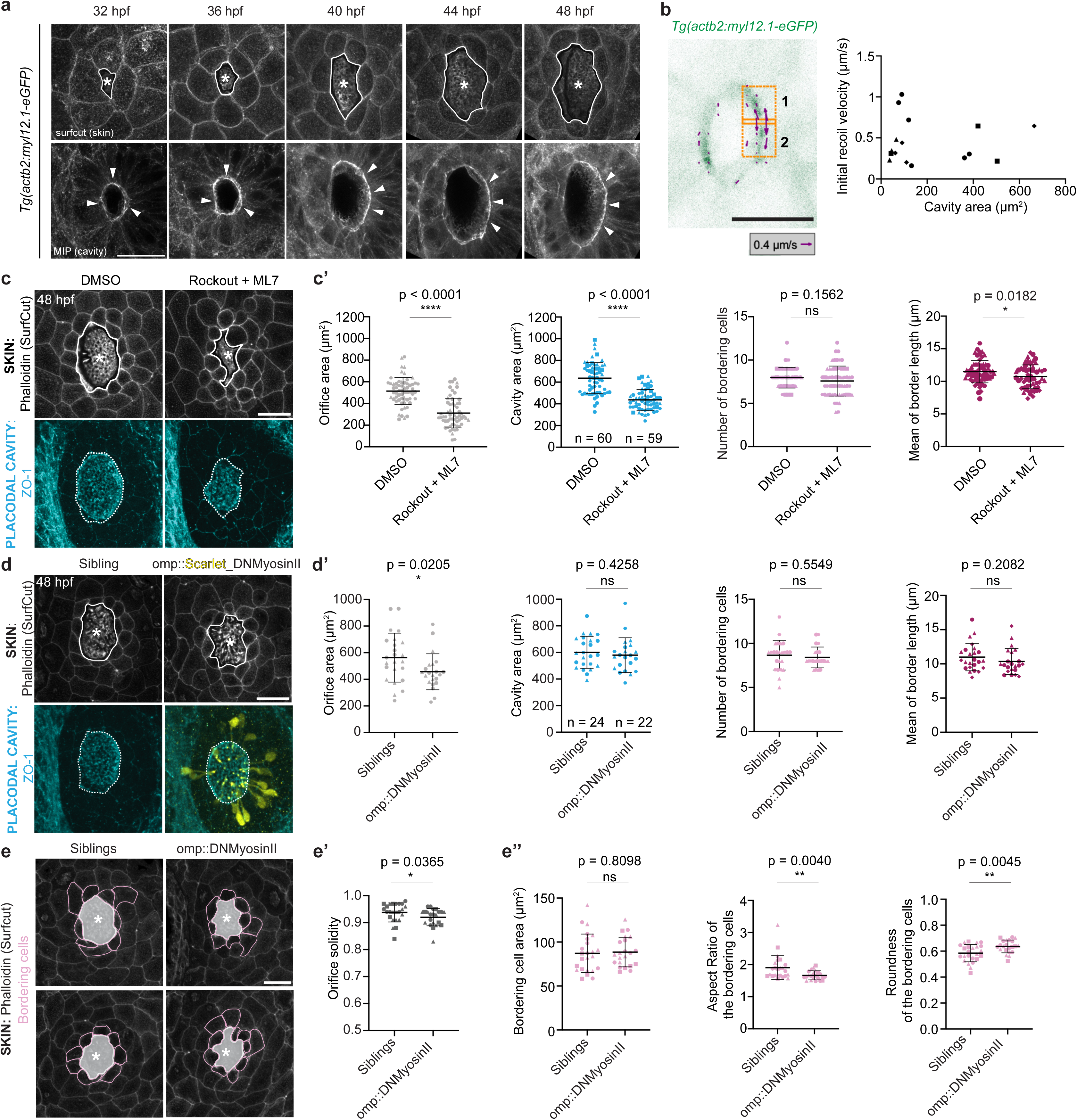
Actomyosin distribution and function. **a.** Time course analysis of non muscle myosin II distribution, visualized with the *Tg(actb2:myl12.1-eGFP)* transgene. Top panels = skin extracted with SurfCut (see Methods). Bottom panels = only the cavity staining is shown (maximum projection), the skin has been removed with the SurfCut plugin. Arrowheads point to the multicellular myosin II ring located in the uppermost part of the cavity. Asterisk = orifice. White line = orifice contour. Scale bar = 20 µm. **b.** Laser ablation of the actomyosin ring. Example of a laser cut performed on the actomyosin ring of a 32 hpf *Tg(actb2:myl12.1-eGFP)* embryo (green). See also Supplementary Movie 7. The image shows the embryo immediately (2s) after the ablation, with tissue velocity field overimposed (purple arrows) in two 5 µm- square boxes on each side of the ablation line. Dorsal view, 1 z-section. Scale bar = 10 µm. The graph shows the initial recoil velocity, used as a proxy for mechanical stress before the cut, as a function of the cavity area (the size of the actomyosin ring was used as a readout of the cavity area, n = 16 ablations, data pooled from 4 independent experiments). **c.** Images of 48 hpf embryos treated with Rockout and ML7 from 32 hpf and DMSO-treated controls. Top images = skin labelled with phalloidin and extracted with the SurfCut plugin (see Methods). Asterisk = orifice. White line = orifice contour. Bottom images = cavity seen with ZO-1 immunostaining, maximum projections. Dotted white line = cavity contour. Scale bar = 20 µm. **c’**. Quantification of orifice area, cavity area, number of bordering cells and mean border length at 48 hpf (4 independent experiments, n = 59 Rockout and ML7-treated embryos, n = 60 DMSO controls). Unpaired, two-tailed t-tests for orifice area and mean of border length; Welch’s t-tests for cavity area and number of bordering cells. **d.** Images of *Tg(ompb:Gal4FF; uas:DNmyosin-Scarlet*) double transgenic embryos at 48 hpf, and control siblings. Top images = skin labelled with phalloidin and extracted with the SurfCut plugin (see Methods). Asterisk = orifice. White line = orifice contour. Bottom images = ZO-1 immunostaining to reveal the cavity (cyan) and Scarlet immunostaining (yellow) to reveal the sensory neurons expressing the dominant negative form of myosin II, maximum projections. Dotted white line = cavity contour. Scale bar = 20 µm. **d’.** Quantification of orifice area, cavity area, number of bordering cells and mean border length (3 independent experiments, n = 22 *Tg(ompb:Gal4FF; uas:DNmyosin-Scarlet*) double transgenic embryos, n = 24 control siblings). Unpaired, two- tailed t-tests for the number of bordering cells and Mann-Whitney tests for the other graphs. **e.** Additional images of the skin (extracted with SurfCut) of *Tg(ompb:Gal4FF; uas:DNmyosin- Scarlet*) embryos and control siblings at 48 hpf, illustrating that the orifice edge is more wiggly in double transgenic embryos and showing the altered shape of the bordering cells. The orifice shape is colored in grey. Scale bar = 20 µm. **e’.** Quantification of orifice solidity. Mann Whitney test. **e’’**. Quantification of bordering cell area, aspect ratio and roundness at 48 hpf (3 independent experiments, n = 22 *Tg(ompb:Gal4FF; uas:DNmyosin-Scarlet*) double transgenic embryos, n = 24 control siblings). Unpaired two tailed t-test for area and roundness, Mann Whitney test for aspect ratio.

Cell divisions orient preferentially along the axis of stretch in many contexts^40^. We hypothesized that tension along the ring plays a non-autonomous role, through physical contact with the overlying skin cells, in triggering or orienting divisions or in regulating skin cell shape. To test this, we used inhibitors of Myosin II phosphorylation, ML7 and Rockout, which target the myosin light chain kinase (MLCK) and the Rho kinase, respectively. When each inhibitor was used individually, no particular phenotype was observed (Supplementary Fig. 3), while combining the two drugs significantly reduced orifice and cavity areas (Fig. 4c, c’). This reveals the implication of the two Myosin II phosphorylation pathways in the coordinated growth of the orifice and placode cavity. The reduced cavity size could be due to decreased cell divisions in the placode and/or impaired intercalation of new cells in the rosette.

To further investigate the role of actomyosin contractility in the placode, we specifically inhibited Myosin II activity in sensory neurons through the mosaic expression of a dominant-negative form of Myosin II. To do so, we combined the *Tg(ompb:Gal4FF)* driver^41^, which starts to be expressed in a subset of ciliated sensory neurons at around 30 hpf (thus preventing any effect on the opening), with the *Tg(uas:DNmyosin-Scarlet)* line^17^. Of note, no Gal4 expression was detected in the MCCs. Interestingly, this perturbation led to a partial uncoupling of the two tissues, with the placodal cavity size remaining unaffected, while the orifice area was significantly reduced compared to siblings (Fig. 4d, d’). The number of bordering cells remained unchanged but they appeared rounder, resulting in a wiggly orifice edge (Fig. 4e, e’, e’’), suggesting that Myosin II activity in rosette neurons influences skin cell morphology, but not the number of their divisions. Overall, these results highlight that actomyosin forces in the placode are important for the smoothening of the growing olfactory orifice in the skin, and for its robust coupling with the cavity.

### ERK signaling is a crucial regulator of orifice expansion and cavity shape

To further investigate the molecular mechanisms involved in orifice expansion, we next focused on the ERK (Extracellular signal-Regulated Kinase) pathway^42,43^, shown to regulate peridermal skin cell proliferation in zebrafish^26^. Immunostaining for phospho-ERK revealed a mosaic signal in subsets of skin cells in the vicinity of the growing orifice, but also further away in the epithelium (Fig. 5a). In the olfactory placode, positive cells with mitotic nuclei were observed outside of the rosette, in basal regions of the placode, and from 44 hpf, a few cells with the shape of sensory neurons were also labeled in the rosette (Fig. 5a’).

**Figure 5.**
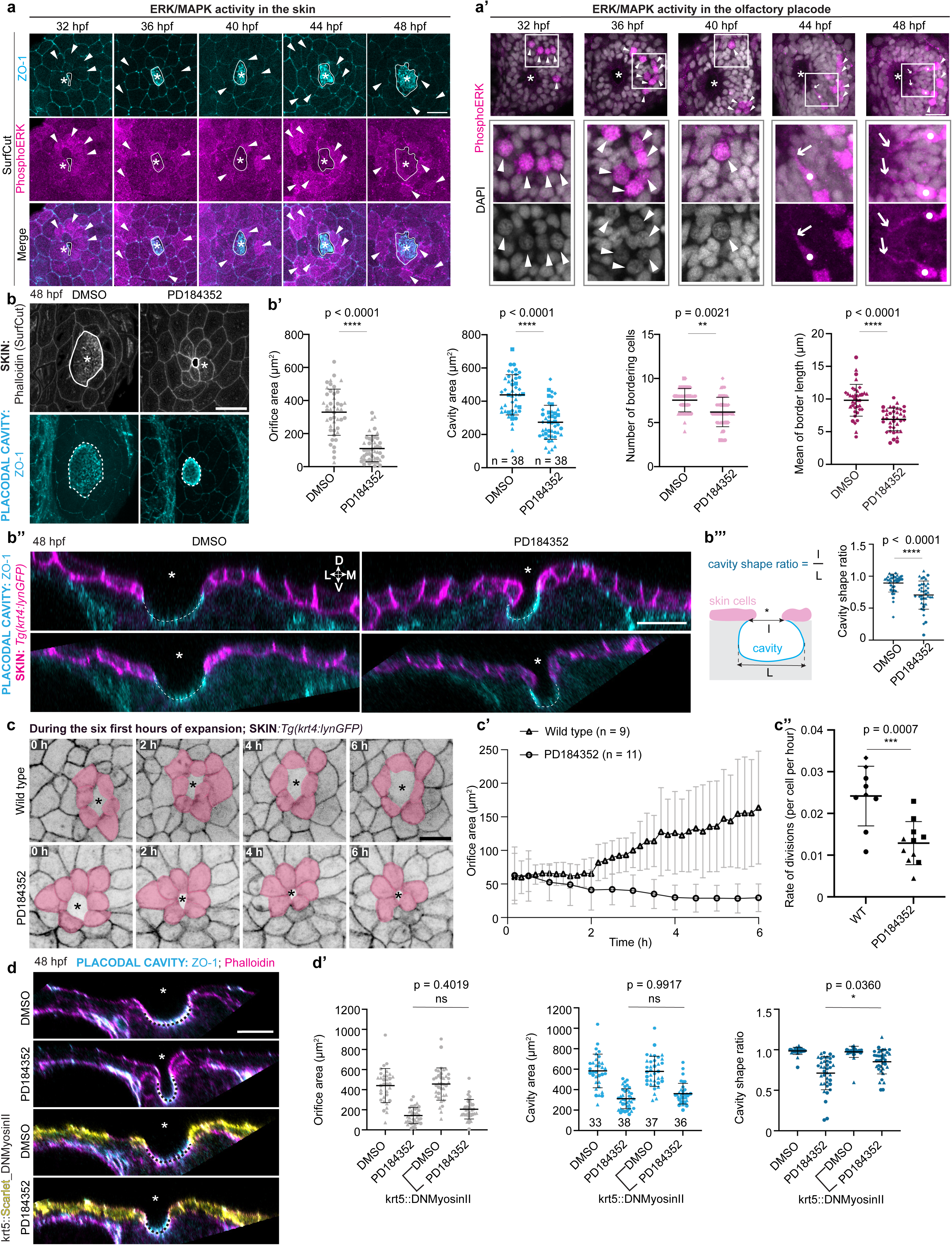
The ERK pathway regulates cavity shape and orifice expansion. **a.** ZO-1 (cyan) and Phospho-ERK (magenta) immunostaining, time course analysis in the skin, extracted with the SurfCut plugin (see Methods). Asterisk = orifice. White line = orifice contour. Arrowheads indicate skin cells showing Phospho-ERK immunoreactivity. Scale bar: 20 µm. **a’**. Phospho- ERK (magenta) immunostaining and DAPI staining (grey), time course analysis in the olfactory placode. Asterisk = cavity. Bottom panels show high magnification views of top panels. Arrowheads point to Phospho-ERK-positive cells located in the basal region of the placode, that appear to be in mitosis based on the DAPI staining. Dots indicate Phospho-ERK-positive placodal rosette cells, and arrowheads point to their dendrites. **b.** Images of 48 hpf embryos treated with PD184352 or DMSO from 32 hpf. Top images = skin labelled with phalloidin and extracted with the SurfCut plugin (see Methods). Asterisk = orifice. White line = orifice contour. Bottom images = cavity seen with ZO-1 immunostaining, maximum projections. Dotted white line = cavity contour. Scale bar = 20 µm. **b’**. Quantification of orifice area, cavity area, number of bordering cells and mean border length at 48 hpf (4 independent experiments, n = 38 PD184352-treated embryos, n = 38 DMSO-treated controls). Mann-Whitney test for orifice area, unpaired two-tailed t-tests for cavity area and number of bordering cells, Welch’s t-test for mean of border length. **b’’**. Orthogonal sections (reslice from the z-stack) of 48 hpf PD184352-treated embryos and DMSO-treated controls carrying the *Tg(krt4:lyn-GFP)* transgene (magenta) to label the skin and immunostained for ZO-1(cyan) to reveal the cavity. Dotted line = cavity contour. Asterisk = orifice. Scale bar = 20 µm. **b’’’**. Quantification of the cavity shape ratio (definition in the schematic view) at 48 hpf (4 independent experiments, n = 38 PD184352-treated embryos, n = 38 DMSO controls). Mann-Whitney test. **c**. Confocal live imaging of the first 6 hours of expansion on *Tg(krt4:lyn-GFP)* wild type embryos or embryos treated with PD184352. Asterisk = orifice. Bordering cells highlighted in pink. See also Supplementary Movie 8. **c’**. Orifice area as a function of time calculated from the movies (2 independent experiments, n = 9 wild type embryos, n = 11 PD184352-treated embryos). Unpaired two-tailed t-tests. **c’’**. Quantification of the division rate in the skin of wild types and PD184352-treated embryos (2 independent experiments, n = 9 wild types, n = 11 PD184352- treated embryos). Unpaired, two-tailed t-tests. **d**. Images of 48 hpf *Tg(krt5:Gal4-ERT; uas:DNmyosin-Scarlet*) double transgenic embryos and control siblings treated with PD184352 or DMSO from 32 hpf. Orthogonal sections (reslice from the z-stack) of embryos immunostained for ZO-1(cyan) to reveal the cavity. Dotted line = cavity contour. Asterisk = orifice. Scale bar = 20 µm. **d’**. Quantification of orifice area, cavity area, and cavity shape ratio upon PD184352 treatment and overexpression of dominant negative myosin II specifically in the skin (3 independent experiments, DMSO on control siblings: n = 33 embryos, PD184352 on control siblings: n = 38 embryos, DMSO treatment on *Tg(krt5:Gal4-ERT; uas:DNmyosin-Scarlet*) embryos: n = 37, PD184352 treatment on *Tg(krt5:Gal4-ERT; uas:DNmyosin-Scarlet*) embryos: n = 36). ANOVA followed by Kruskal-Wallis multicomparison test.

To test the role of ERK signaling, we pharmacologically inhibited the pathway from 32 hpf using PD184352, an inhibitor of the ERK upstream regulator, MEK1/2. The efficiency of the inhibition was visualized by the loss of phospho-ERK immunoreactivity in the skin and placode (Supplementary Fig. 4a). This led to a strong phenotype at 48 hpf, with orifices that were much smaller than in DMSO-treated controls, as well as a reduced number of bordering cells and border length (Fig. 5b, b’). Live imaging of PD184352-treated embryos during the first 6 hours of expansion (progressive drifts of the embryos along the Z-axis prevented us to image the full initial expansion phase in this condition) showed delayed growth and in some cases a shrinkage of the olfactory orifice over time, leading to its partial or complete closure (Fig. 5c, c’ and Supplementary Movie 8). The number of skin cell divisions was reduced in our movies (Fig. 5c’’), consistent with the previously described role of ERK signaling in peridermal skin cell proliferation^26^. The rosette cavity was also smaller in treated embryos (Fig. 5b, b’), indicating a potential role for ERK in placode cell divisions or addition of new cells into the rosette.

Strikingly, close examination of orthogonal sections revealed an abnormal purse-like shape of the cavities in PD184352-treated embryos at 48 hpf, with the upper, apical part of the cavity being smaller than its basal region (Fig. 5b’’, b’’’). This cavity deformation was not observed upon HUA-mediated inhibition of cell divisions (Supplementary Fig. 4b). This suggests that, in addition to its impact on cell divisions, ERK signaling inhibition induces an aberrant distribution of mechanical forces resulting in the constriction of the upper cavity. To test whether these aberrant forces are generated in the skin, we overexpressed the dominant-negative form of Myosin II specifically in the skin in PD184352 and DMSO-treated embryos, using the *Tg(uas:DNmyosin-Scarlet*) and *Tg(krt5:Gal4-ERT)* transgenes. While this did not rescue the cavity and orifice sizes, it significantly rescued the purse-like shape of the cavity (Fig. 5d, d’). This result strongly suggests that ERK inhibition increases actomyosin contractility in skin cells, which contributes to the constriction of the upper part of the placodal cavity, leading to its abnormal shape. This points to a role of ERK-mediated relaxation in the skin epithelium for maintaining the open shape of the placode cavity during orifice expansion, thus revealing a mechanical feedback from the skin back to the olfactory placode rosette.

## Discussion

In this study, we characterized the cell behaviors and reciprocal tissue interactions that guide the growth and morphogenesis of the zebrafish nostril orifice during the first 16 hours following its opening. Our analysis highlights the dynamic rearrangements of the surrounding skin epithelium, including cell deformations and divisions - though the latter are dispensable for orifice growth. We propose a model (Fig. 6a) in which cavity growth, occurring through gradual incorporation of new placode cells in the rosette, generates outward forces that are transmitted to the skin through cell adhesion, allowing the border of the placodal cavity to act as a template for epithelial hole enlargement. In addition to this main scaling mechanism, a neuronal actomyosin ring formed in the upper cavity further adjusts the tissue-tissue coupling by smoothening the edge of the epithelial orifice. Last, ERK-mediated epithelial relaxation is required in the skin to keep the open shape of the placodal cavity during orifice growth, demonstrating a reciprocal mechanical interplay between the two tissues (Fig. 6b).

**Figure 6.**
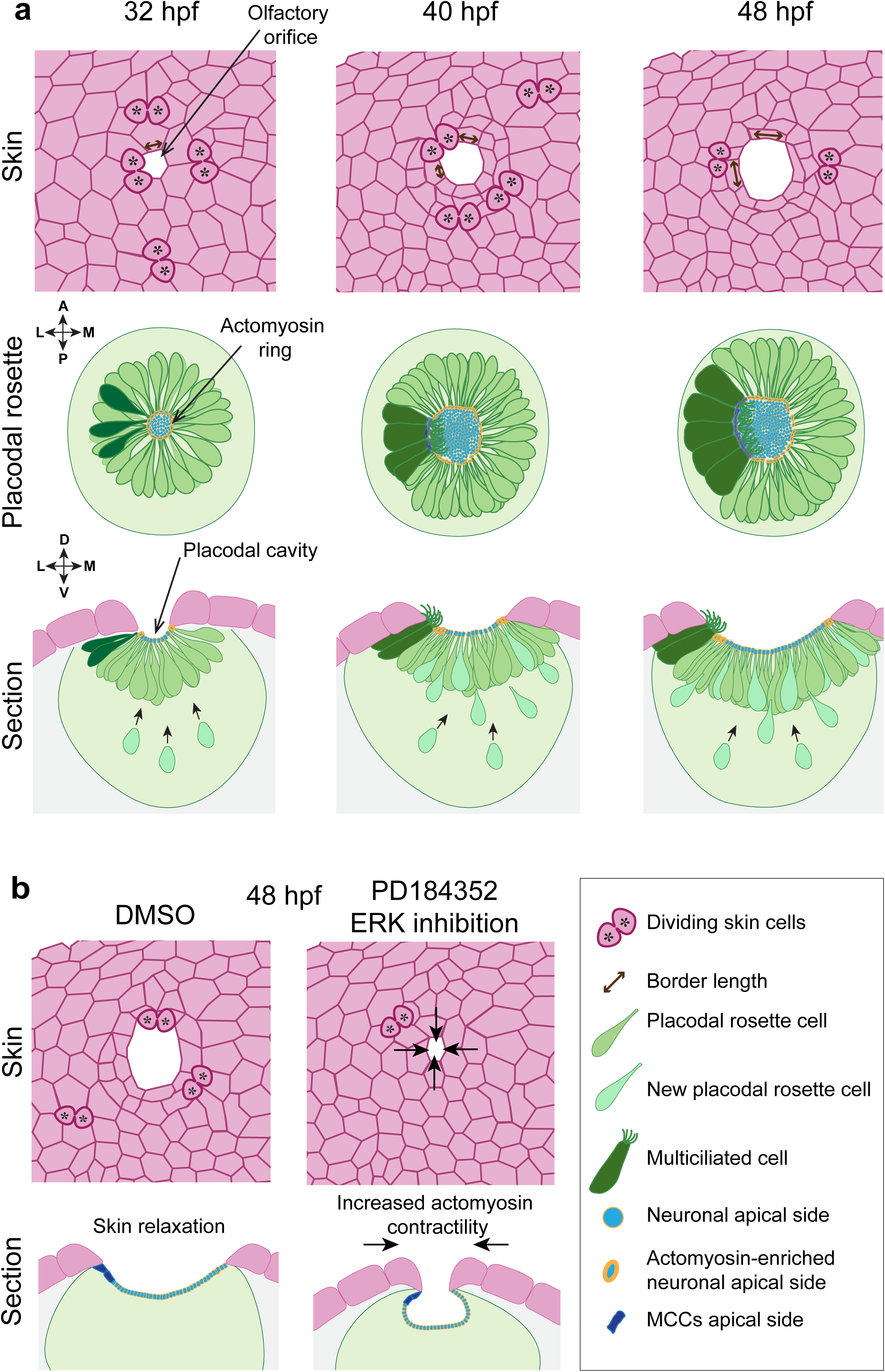
Proposed model for the expansion and shaping of the olfactory orifice. **a.** Schematic view of the cellular processes and mechanisms contributing to the expansion and shaping of the olfactory orifice at 32, 40 and 48 hpf. Top row, dorsal view of the skin epithelium, skin cells near the orifice exhibit increased cell divisions (marked by asterisks) which align with the edge but are dispensable for orifice growth. Bordering skin cells also undergo cell shape changes: they become less round/more elongated and increase their border length (indicated with double arrows). Middle row, dorsal view of the underlying olfactory placode. At 32 hpf, the placode forms a rosette structure, primarily composed of sensory neurons (light green) and a few undifferentiated MCCs (dark green), whose apical tips (light and dark blue, respectively) cluster to form a central placodal cavity. The number of apical surfaces increases over time due to intercalation of new placodal cells (see third row). Cortical actomyosin is present around the small, round apical tips and is particularly enriched at the interface between the skin and the cavity, which results in the formation of a multicellular actomyosin ring. Third row, cross section of the placode and skin tissues. New cells coming from more basal regions of the placode (lightest green) continuously join the rosette, extending the cavity. Concomitantly, MCCs differentiate in the lateral cavity, which also participates in cavity expansion. The progressive widening of the cavity produces forces that are transmitted to the skin through adhesion, leading to a scaled orifice expansion. In addition, the multicellular actomyosin ring acts non-autonomously on the skin epithelium to smoothen the orifice margin, reinforcing the alignment and coupling between the two tissues. **b.** Schematic view of the consequences of ERK inhibition on orifice expansion analyzed at 48 hpf. First row, inhibition of ERK signaling disrupts orifice expansion, resulting in its partial collapse (arrows). Embryos treated with the ERK inhibitor show reduced skin cell proliferation. Second row, the cavity is smaller and acquires an abnormal purse-like shape upon ERK inhibition, which is at least partially due to an increase in actomyosin contractility in skin cells. This reveals a role for skin relaxation to maintain the open cavity shape in the control situation, and supports the idea of a bi-directional mechanical coupling between the two tissues.

We described an enrichment of skin cell divisions close to the growing orifice, with their division axis aligning with the edge of the hole. Unexpectedly, impairing skin cell divisions did not affect the growth of the olfactory orifice, which occurred through drastic shape changes of the bordering cells. Even though cell divisions are not essential for the orifice to reach its proper size at this stage, providing new skin cells on the orifice edge through cell division may prevent excessive cell deformation and further epithelial rupture. This could also be important for later stages of expansion^27,44^ and remodeling of the orifice/cavity structure or the future nostril homeostasis. In addition, cell divisions could promote tissue fluidity in this area^21,22,45^ or buffer the forces emanating from the cavity/orifice border^46^. What is the signal regulating the rate and orientation of cell divisions around the orifice? The number of bordering cells was only slightly decreased in conditions where the placode/cavity growth is impaired (laser ablation and *gmnc* mutant). These placode-specific perturbations may not be drastic enough to see a clear effect on the number of bordering skin cells, leaving open the question of the role of the placode in regulating skin cell divisions. ERK signaling seems important to reach a proper rate of peridermal skin cell divisions, consistent with previous studies in the zebrafish tail^26^. The partial loss of adhesion with surrounding neighbors by skin cells located at the orifice edge (i.e. the presence of free space) could promote cell divisions locally, as cell/cell contacts are known to inhibit cell proliferation in other contexts^47^. Additional experiments will be required to further understand the function and regulation of skin cell divisions around the orifice.

Our results reveal that the lengthening of the skin cell membranes along the orifice edge is an important contributor to the growth process. Similar cell shape changes have been described to occur on the edge of epithelial orifices *in vitro*^15,16^, and tangential elongation of the apical domain through actin polymerization allows initial lumen expansion in a human epiblast model^48^. Epithelial thinning, another type of cell shape change, is also important for lumen growth in the zebrafish otic placode^23^. The precise molecular mechanism by which the skin cells change their shape and devote more and more membrane to the edge of the orifice remains to be clarified, and could involve actin polymerization, membrane addition pathways and/or adhesion molecules to establish new junctions with the apical areas of the placodal rosette cells lining the underlying cavity.

We identified the growth of the cavity in the center of the placodal rosette as an essential driver of orifice expansion. We showed that cavity growth results from the progressive addition of new apical areas over time. We mentioned the presence of sensory neurons and MCCs in the rosette, but additional cell types may also contribute to forming the rosette cavity at these stages, including olfactory rod cells^44^ and ionocytes^49^. We propose that the outward forces produced by the cavity growth are transmitted to the orifice edge through tight-junction-mediated adhesion between the two tissues, occurring in the upper part of the cavity. This process would allow the epithelial hole to grow in a way that matches the size of the underlying cavity and placodal tissue. In other words, the rosette cavity would serve as a physical scaffold for the growth of the epithelial hole.

Interestingly, the coordination between the growth of the placodal rosette and the epithelial orifice is further fine-tuned by the multicellular actomyosin ring located in the uppermost placodal cavity. Our results show that this ring is under tension and has a non-autonomous role in regulating skin cell morphology to smoothen the orifice edge, while no influence on the number of skin cell divisions was detected. An actomyosin ring has been described around the holes forming spontaneously *in vitro* in epithelioid tissues of human embryonic stem (hES) cells, where it is also involved in the smoothening the orifice margin upon local ruptures^9^, but the ring is located within epithelial cells and thus acts in a tissue-autonomous manner in that context. Multicellular actomyosin rings usually are contractile machines actively closing epithelial holes during embryonic development, wound healing or apoptotic cell extrusion^50–52^. However, in our system the ring-like actomyosin structure coexists with the growth of the rosette cavity and the expansion of the epithelial hole. How the tensile forces along the ring are finely regulated and/or balanced by other force-generating processes to allow the expansion of the cavity and olfactory orifice remains to be elucidated.

Pharmacological inhibition of ERK signaling led to a severe nostril phenotype, with small cavity and orifice, pointing to this pathway as a key player in the expansion of the olfactory orifice. The number of cell divisions was reduced in the skin in this condition. However, since skin cell divisions are dispensable for orifice growth, this is unlikely to contribute to this drastic defect. The rosette cavity was also smaller in treated embryos, suggesting a role for ERK in placode cell divisions or cell intercalation within the rosette.

On top of these effects on skin cell division and cavity size, ERK inhibition resulted in a novel phenotype: the abnormal purse-like shape of the cavity, with a constriction in its uppermost region. Indeed this cavity shape defect was not seen upon HUA-mediated inhibition of proliferation, suggesting that it is due to a mechanism that is distinct from the reduction in cell divisions. Strikingly, this cavity shape phenotype was partially rescued upon skin-specific perturbation of Myosin II activity. This suggests that in the wild type situation ERK signaling downregulates actomyosin contractility in the skin epithelium, which would maintain the open shape of the rosette cavity and promote orifice growth.

Links between cell mechanics and ERK have been previously described, in which ERK signaling acts as a mechanotransduction pathway downstream of mechanical cues^53^ or, as it is the case in our system, acts upstream of the actomyosin cytoskeleton^43^. For example, ERK has been proposed to activate the contractile machinery during cell migration through the regulation of myosin light chain kinase (MLCK) and myosin phosphatase activities^54,55^. In addition, ERK is known to increase actin polymerization at the edge of protrusions in migrating cells^56,57^. Future experiments will investigate which mechanism is at play downstream of ERK signaling to decrease skin contractility. Interestingly, tissue relaxation through a reduction in actomyosin activity has also been proposed to participate in tissue morphogenesis in other contexts. For instance, local downregulation of actomyosin at tricellular junctions is required for the transient opening of the follicle epithelium during Drosophila oogenesis^6^, and at a larger scale, epithelial relaxation has been described during ventricle lumen expansion in the zebrafish hindbrain^58^, zebrafish epiboly and tissue elongation in Drosophila^59^.

Overall this work uncovers a previously unrecognized tissue-tissue coupling mechanism that coordinates epithelial hole expansion with the growth of a sensory placode-derived cavity, and illustrates more broadly how reciprocal interactions between adjacent tissues sculpt organ morphogenesis.

## Materials and methods

### Zebrafish lines

Wild-type and transgenic zebrafish embryos were obtained by natural spawning. In the text, the developmental times in hpf indicate hours post-fertilization at 28.5°C. For all stages embryos were collected at 10 am. To obtain the 36 hpf stage, embryos were incubated at 28.5°C for 8h before being placed at 33°C overnight, followed by incubation at 28.5°C until fixation time at 6 pm. For the stages 40 and 44 hpf, embryos were incubated at 28.5°C until 28 hpf and then placed at 25°C overnight. The next morning they were placed back at 28.5°C and fixed at 10 am for 40 hpf and at 2 pm for 44 hpf.

We used the following published lines (the simplified names are used in the figures and their legends): *Tg(krt4:LY-eGFP)^sq^*^18^ (reference^19^, a gift from Tom Carney, Singapore, referred to as *Tg(krt4:lynGFP)*), and *Tg(krt5:Gal4-ERT2-VP16, myl7:CFP)^b1234^* (reference^60^, referred to as *Tg(krt5:Gal4-ERT)*), were used to label and target the peridermal skin cells; *Tg(- 8.0cldnb:LY-EGFP)^zf106^* (reference^61^, referred to as *Tg(cldnb:meGFP)*) was used to visualize both peridermal and placodal cells; *Tg(-2.0ompb:gapYFP)^rw032^* (reference^30^, a gift from Nobuhiko Miyasaka, RIKEN Institute, National Bioresource Project of Japan, referred to as *Tg(omp:meYFP)*) and *Tg(Xla.Tubb:DsRed)^hkz^*^018t^ (reference^62^) were used to visualize olfactory placode neurons; *Tg(14Xuas:mRFP, Xla.Cryg:GFP)^tpl^*^2^ (reference^63^, referred to as *Tg(uas:RFP)*), was used to label the cytoplasm of Gal4-expressing cells; *Tg(actb2:myl12.1- eGFP)^e2212^*, referred to as *Tg(actb2:myl12.1-eGFP),* was used as a reporter line to visualize non muscle myosin II^37^. *Tg(uas:DNmyosin-Scarlet)^bps^*^5^ was used to inhibit myosin II activity^17^.

Two lines were produced for this study. The *Tg(uas:cdkn1b-E2A-GFP)^bps6^* line was generated by Tol2-mediated transgenesis (see plasmid construction) to inhibit cell divisions in a tissue-specific manner. In order to drive specific UAS lines in the tissues targeted by the enhancer-trap line *Et(krt4:EGFP)^sqet^*^33^ from reference^25^, we performed a CRISPR-driven knock-in of a E2A-KalTA4 sequence into the coding sequence of eGFP, as described in reference^64^. Accordingly, the resulting *Tg(KALTA4)^bps7^* line, referred to as *Tg(krt4:KalTA4),* drives a robust expression in peridermal skin cells and roof plate cells.

*gmnc^sq^*^34^ mutants^32^, *dnaaf1^tm^*^317b^ mutants^34,35^ and *traf3ip1^tp^*^49d^*/elipsa* mutants^34,36,65^ were used to analyze the role of MCCs and cilia.

All our experiments were made in agreement with the European Directive 210/63/EU on the protection of animals used for scientific purposes, and the French application decree ‘‘Décret 2013–118’’. The fish facility has been approved by the French ‘‘Service for animal protection and health’’, with the approval number B-75-05-25. The DAP number of the project is APAFIS #47427-2023121817395554 v6.

### Plasmid construction

The Tol2 *uas:cdkn1b-E2A-meGFP* plasmid was designed by the authors and synthesized by Twist Bioscience. The coding sequence for the degradation-resistant mutant of human CDKN1B (cyclin-dependent kinase inhibitor 1B)^24^ was optimized for zebrafish expression using iCodon^66^. It was fused to E2A peptide and meGFP coding sequences, and placed under the control of 5XUAS.

### Immunostainings

For immunostaining, embryos were fixed in 4% paraformaldehyde (PFA, in PBS) overnight at 4°C or 2 hours at room temperature, blocked in 3% goat serum and 0.3% triton in PBS for 2h at room temperature or overnight at 4°C and incubated overnight at 4°C with primary and secondary antibodies. The following primary antibodies were used: anti-GFP (chicken, 1/200, GFP-1020, Aves labs), anti-DsRed (rabbit, 1/300, 632496, Takara Bio), anti- ZO1 (mouse IgG1, 1/500, 33-9100, ThermoFisher scientific), anti-Centrin (mouse IgG2a, 1/200, 04-162, Millipore), anti-Acetylated tubulin (mouse IgG2b, 1/500, T67973, Sigma), and anti-PhosphoERK (rabbit, 1/200, 4370, Cell Signaling). For phalloidin staining, an overnight incubation was performed at 4°C with phalloidin-rhodamine (1/200, R415, ThermoFisher scientific) or phalloidin-Alexa488 (1/200, A12379, ThermoFisher scientific).

### Drug treatments

Embryos were dechorionated and incubated from 32 hpf to 48 hpf in four-well plates with the drugs or the equivalent % of DMSO diluted in E3 medium. To inhibit proliferation, embryos were treated with 20 mM hydroxyurea and 150 µM aphidicolin (Sigma, A0781) (HUA). To block Myosin II phosphorylation, embryos were incubated in 50 µM Rockout (Calbiochem, 555553) and 50 µM ML-7 (Sigma, I2764), alone or in combination. PD184352 (Sigma, PZ0181) treatment at 10 µM was used to inhibit MEK1/2 (Sigma, PZ0181). To induce the expression of UAS transgenes with the *Tg(krt5:Gal4-ERT)* line, the embryos were treated at 9 hpf with 4-hydroxy-tamoxifen (Sigma, H7904) dissolved in ethanol (final concentration of 8 ng/mL).

### Image acquisition

For live imaging, embryos were dechorionated, tricained (0.016%) and mounted at 32 hpf in 0.5% low melting agarose in 1X E3 medium. Movies were recorded at 28.5°C on a Leica TCS SP8 MPII upright multiphoton microscope using a 63X (NA 0.9) water lens. For fixed embryos, immunostained embryos were mounted in 0.5% low melting agarose in PBS and imaged on a Zeiss 980 FAST Airyscan with a 20X (NA 1.0) water lens.

### Laser ablations

For ablating cells of the placodal rosette, embryos at 32 hpf were dechorionated, tricained (0.016%) and mounted in 0.5% low melting agarose in 1X E3 medium and imaged using a Leica TCS SP8 MPII upright multiphoton microscope using a 25X (NA 0.95) water lens. Ablations were performed with a pulsed femto-second laser tuned at 790 nm. A z-stack was acquired before and after each ablation. After ablation, embryos were kept mounted in 0.5% low melting agarose in 1X E3 medium overnight at 28.5°C and fixed for immunostaining analysis at 48 hpf.

For the ablation of the actomyosin ring, embryos were dechorionated, tricained (0.016%) and mounted in 0.5% low melting agarose in 1X E3 medium, on the bottom of Ibidi dishes (81158). The dishes were flipped and imaged using a Leica TCS SP8 MPII upright multiphoton microscope using a 63X oil objective (NA 1.4). Ablations were performed using a pulsed femto-second laser tuned at 790 nm. Only the movies in which a clear cut could be observed (i.e. no fluorescence recovery overtime) were kept for further analysis. To analyze the recoil, we measured the local flow by particle image velocimetry using the MatPIV toolbox for Matlab (Mathworks, US). The window size was set to 32 pixels (between 2.5 and 5 µm depending on the developmental stage), with an overlap of 0.5. The kinetics of relaxation was assessed between the pre-cut time-point and 1 frame later (around 2s) in two 5 µm-square boxes on each side of the region of ablation, indicated by the squares in Fig. 4b. The “recoil velocity” was then obtained with the following formula: Vrecoil = <V>2 - <V>1 where V indicates the average velocity component orthogonal to the cut line, and < > denotes the average over the 5 µm-square boxes (1 or 2, see Fig. 4b), as previously performed in references^17,67^.

### Image analysis and quantifications

#### Analysis of skin cells, olfactory orifice and placodal cavity in fixed samples

In fixed embryos in which peridermal skin cells were labelled with membrane or cytoplasmic fluorescent proteins or with phalloidin staining, the most superficial layer was isolated using the SurfCut plugin^68,69^. This plugin allows to extract and project, from a 3D sample with a slightly curved geometry (which is the case of the skin overlying the placode), the fluorescent signal coming exclusively from the sample surface. It also allows to extract the signal of a 3D sample at a given distance from the sample surface, which can be chosen by the user. The area of the olfactory orifice was measured on 2D maximal projections, by outlining the olfactory orifice with the freehand selection tool and ROI manager plugin of ImageJ. The number of bordering cells was analyzed by counting the number of skin cells directly lining the orifice, on 2D maximal projections. The mean border length was obtained by averaging the length of the cell membrane portion facing the orifice, which was measured with the freehand line tool on ImageJ. ZO-1 immunostaining or phalloidin staining were used to analyze the area of the placodal cavity through the freehand selection tool and ROI manager plugin of ImageJ, on 2D maximal projections. This step was performed in a blind manner, meaning without visualizing the overlying orifice in the skin. The solidity of the orifice edge was obtained by dividing the orifice area by its convex hull area. Bordering cells were segmented with the SAMJ-IJ plugin (https://github.com/segment-anything-models-java/SAMJ-IJ). Their aspect ratio was then obtained by fitting an ellipse to the cell and calculating the major axis / minor axis ratio of the ellipse. The roundness of the bordering cells were calculated as follows: (4 x (area))/(pi x major axis^2^).

#### Skin segmentation

To describe and quantify the skin remodeling events in our movies (Fig. 2), the contours of skin cells and the orifice were semi-automatically segmented with the Python version of Tissue Analyzer (pyTA), using the skin membrane GFP labelling in the *Tg(krt4:lynGFP)* line. The same program was used to manually correct the segmentation and to perform cell tracking. The first and second neighbors were manually annotated on the last frame of the movies, and backtracked using a Matlab (Mathworks, US) custom program. This program was also used to measure the orifice area and perimeter over time, and to analyze the average border length, by tracking the “junction” between the bordering cells and the orifice over time. To determine the bordering cell area or aspect ratio (Supplementary Fig. 1b, b’ and Fig. 4e, e’’), the SAMJ-IJ plugin was used.

#### Skin cell divisions

To obtain the map of skin cell divisions for each embryo, the positions (x,y) of the two daughter cells after division were manually annotated using the ImageJ plugin MaMuT^70^. This information was then processed by a custom Matlab (Mathworks, US) program to generate the division maps in Fig. 2 and Supplementary Fig. 1. On these maps the dots indicate the positions of the two daughter cells, which are linked by a straight segment. To assess the orientation of cell divisions with respect to the edge of the orifice, the orifice was first fitted with an ellipse. The absolute value of the smallest angle (between 0° and 90°) that the axis of division forms with the tangent to the ellipse was then calculated, as depicted in Fig. 2c’’’. The rate of cell divisions was obtained by dividing the number of cell divisions in a given cell category (first/second neighbours and rest of the epithelium) by the number of cells of this category at the beginning of the movies and by the duration of the movies.

#### Segmentation of the apical surfaces of placodal cells lining the cavity

The deep learning based tool StarDist^71^ was used to detect and segment the contours of the small, round apical sides lining the placodal cavity revealed by ZO1 immunostaining. The contours of the large apical surfaces of MCCs were manually segmented following a 3D reconstruction in Arivis.

### Statistical analysis

Graphs show means ± standard deviation, overlayed with all individual data points. The plots were generated with the GraphPad Prism software. On the plots, embryos from the same experiment are represented with similar markers (dots, triangles, squares, etc). The nature of the statistical test performed for each graph is indicated in the figure legends. For all graphs, we checked for normality of the data distribution and analyzed the variance before choosing to perform parametric, unpaired, two tailed t tests (or ANOVA tests for multiple comparisons) or adequate non parametric tests. To compare the distribution of angles in Fig. 2c’’’, we first evaluated the uniformity of the angle distributions. We computed the lengths of the resultant vectors for the two angle distributions (R1 and R2): the greater the value of R (close to 1), the more the angles are concentrated around one direction. Then, we compared the 2 concentrations (peakedness κ) of the 2 distributions using a permutation test (bootstrap). The p values correspond to *p < 0.05, **p < 0.01, ***p < 0.001.

## Acknowledgements

We acknowledge Christine Vesque and Alain Trembleau for their insightful comments in reviewing the manuscript. We are grateful to Nathalie Jurisch-Yaksi and Sudipto Roy for providing us with *gmnc* mutant embryos and *gmnc* mutant fish, respectively. This work was funded by the Agence Nationale pour la Recherche (ANR-23-CE13-0025 MECAMATRIX), the Centre National pour la Recherche Scientifique (CNRS), Sorbonne Université (including a grant from the i-Bio initiative), and the National Institute of Health NIDCD Grant R01-DC- 017989. C. Gordillo Pi was supported by a doctoral fellowship from the “Ministère de l’Enseignement supérieur et de la Recherche” and a 4th year fellowship from the Fondation ARC pour la recherche sur le cancer (ARCDOC42024010007666). We also thank the imaging platform of the Institut de Biologie Paris-Seine (the facility is supported by CNRS, Sorbonne Université and the Conseil Régional Ile-de-France) and the IBPS aquatic platform for fish care.

