## Supplementary_Figures for "Dynamic interactions between epithelial skin cells and a sensory cavity sculpt the growing olfactory orifice"

Supplementary Figure 1 Gordillo Pi et al.

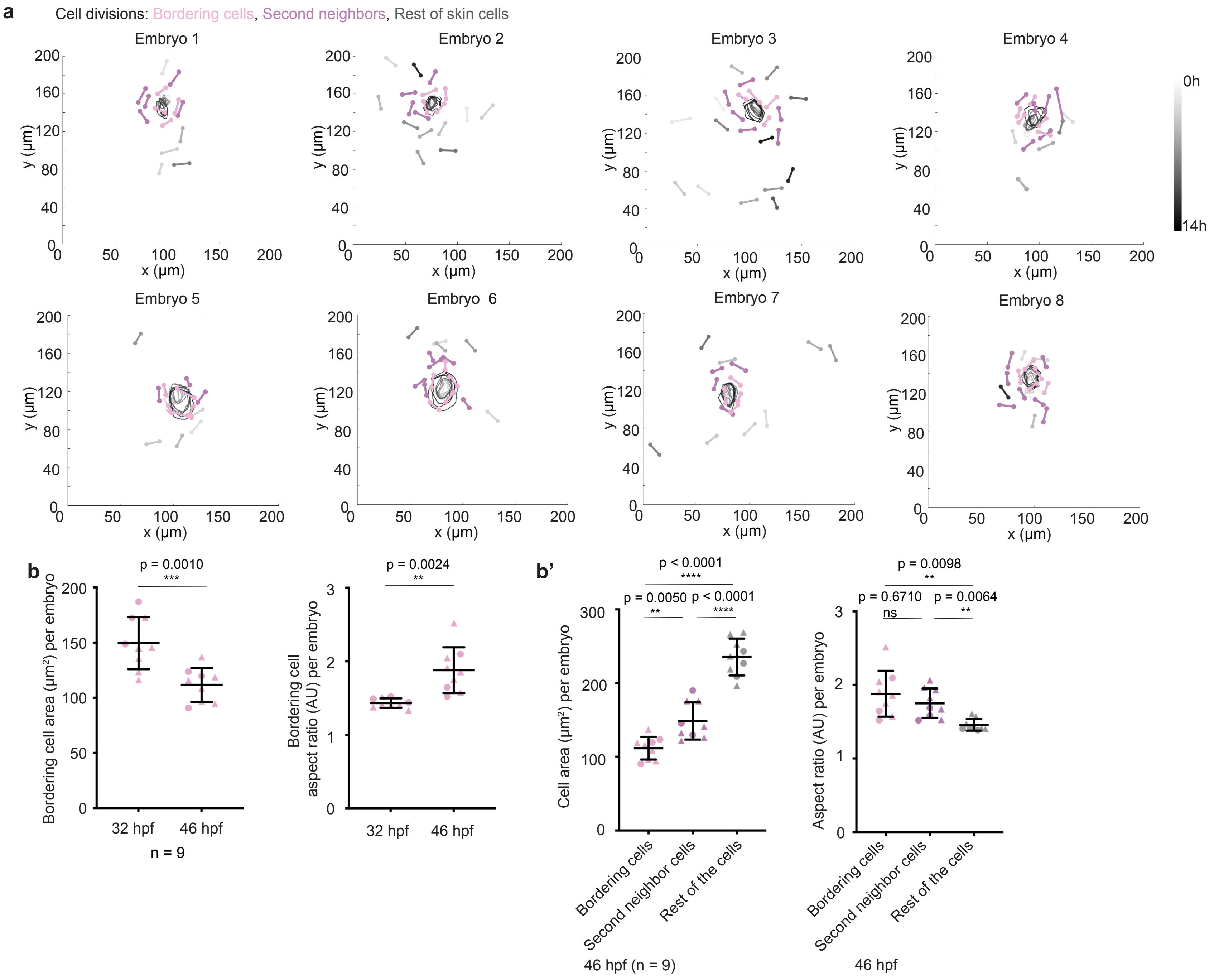

**Supplementary Figure 1. Additional maps of cell divisions and quantification of skin cell remodeling during orifice expansion,** related to Fig. 2**. a.** Map of cell divisions occurring during the time lapse sequence in the field of view, related to Fig. 2c'. Grey lines = contours of the growing orifice, the time is color coded. For each division, a segment links the centers of the two daughter cells, which are represented by the dots. Pink segments = 1st neighbor skin cell divisions, purple segments = 2nd neighbor cell divisions, grey segments = divisions in the rest of the skin epithelium (for those, time is color coded). **b.** Comparison of the area or aspect ratio of bordering cells at 32 and 46 hpf (n = 9 embryos from 2 independent experiments). Unpaired, two-tailed t-test for the area, and Welch's t-test for the aspect ratio. **b'.** Comparison of the area or aspect ratio of bordering cells, second neighbor cells and rest of the skin cells at 46 hpf (9 embryos from 2 independent experiments). ANOVA followed by Tukey multiple comparison test for the cell area, Welch's ANOVA followed by Dunnett T3 multiple comparison test for the aspect ratio.

Supplementary Figure 2 Gordillo Pi et al.

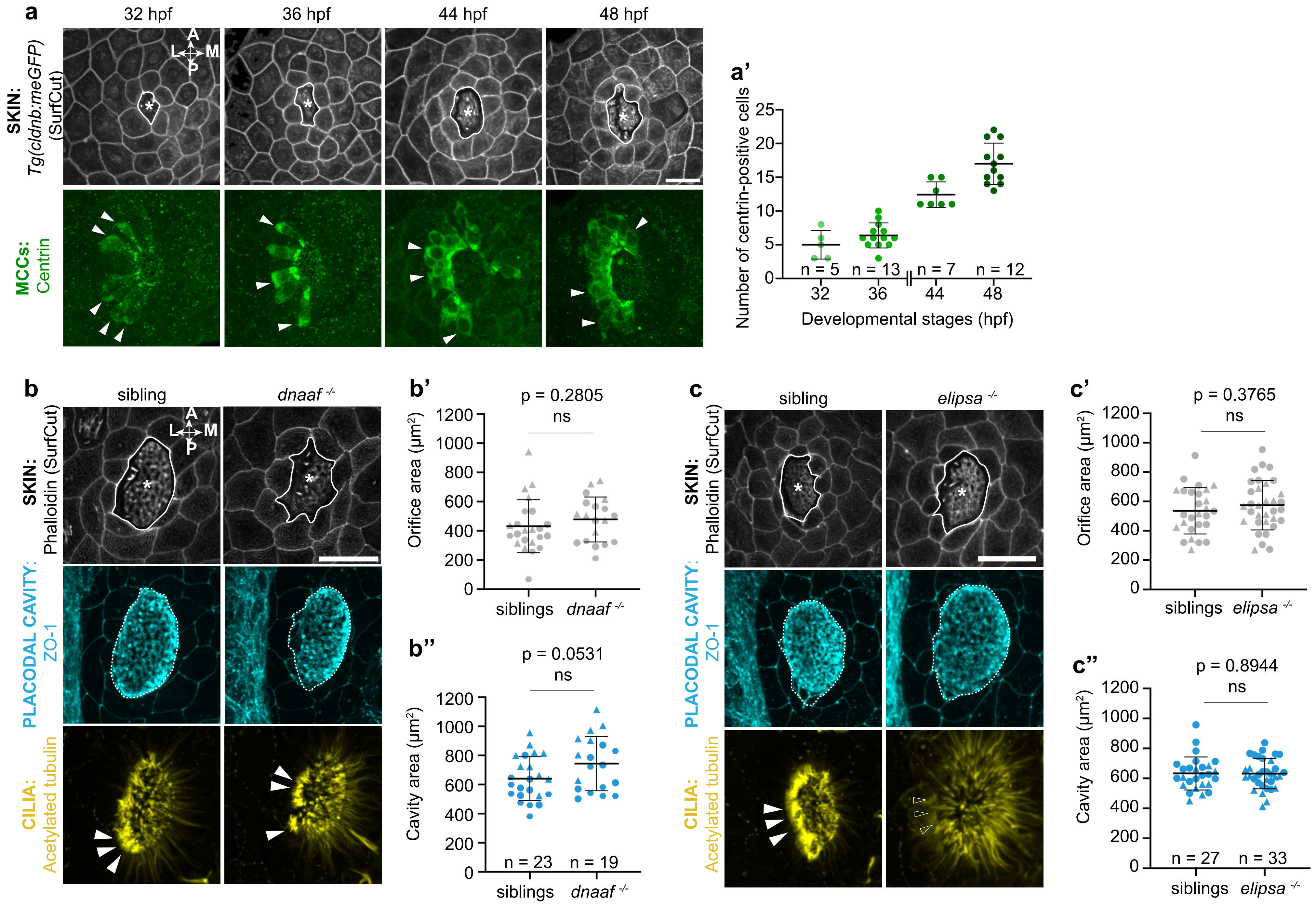

**Supplementary Figure 2. Additional data on MCCs and analysis of ciliary mutants,** related to Fig. 3**. a.** Images of *Tg(cldnb:meGFP)* embryos immunostained for Centrin, time course analysis. Top panels show *Tg(cldnb:meGFP)* expression in the skin (grey) extracted with the SurfCut plugin, bottom panels show Centrin expression (green, 1 z-section). Arrowheads point to Centrin-positive MCCs on the lateral side of the cavity. Dorsal view. Asterisk = orifice. White line = orifice contour. Scale bar: 20 µm. **a'.** Number of Centrine-positive cells, time course analysis (1 experiment, 32 hpf: n = 5 embryos; 36 hpf: n = 13 embryos; 44 hpf: n = 7 embryos; 48 hpf: n = 12 embryos). **b.** Images of *dnaaf* $^{-/-}$ mutant embryos and control siblings at 48 hpf. Top images = skin labelled with phalloidin (grey) and extracted with the SurfCut plugin, middle images = cavities seen with ZO-1 immunostaining (maximum projections), bottom images = Acetylated-tubulin immunostaining to reveal the cilia (more generally acetylated microtubules). Asterisk = orifice. White line = orifice contour. Dotted white line = cavity contour. Arrowheads indicate the multiple cilia of MCCs in the lateral region of the cavity, present in both controls and *dnaaf* $^{-/-}$ mutants. Scale bar = 20 µm. **b', b''**. Quantification of orifice area (b') and cavity area (b'') in *dnaaf* $^{-/-}$ mutants and control siblings at 48 hpf (3 independent experiments, n = 19 *dnaaf* $^{-/-}$ mutants, n = 23 control siblings). Mann-Whitney test for orifice area, and unpaired, two-tailed t-test for cavity area. **c.** Images of *elipsa* $^{-/-}$ mutant embryos and control siblings at 48 hpf. Top images = skin labelled with phalloidin (grey) and extracted with the SurfCut plugin, middle images = cavities seen with ZO-1 immunostaining (maximum projections), bottom images = Acetylated-tubulin immunostaining to reveal the cilia (more generally acetylated microtubules). Asterisk = orifice. White line = orifice contour. Dotted white line = cavity contour. Arrowheads indicate the multiple cilia of MCCs in the lateral region of the cavity, present in control siblings but not in *elipsa* $^{-/-}$ mutants. Scale bar = 20 µm. **c', c''**. Quantification of orifice area (c') and cavity area (c'') in *elipsa* $^{-/-}$ mutants and control siblings at 48 hpf (2 independent experiments, n = 33 *elipsa* $^{-/-}$ mutants, n = 27 control siblings). Unpaired, two-tailed t-test for orifice area, and Mann-Whitney test for cavity area.

Supplementary Figure 3 Gordillo Pi et al.

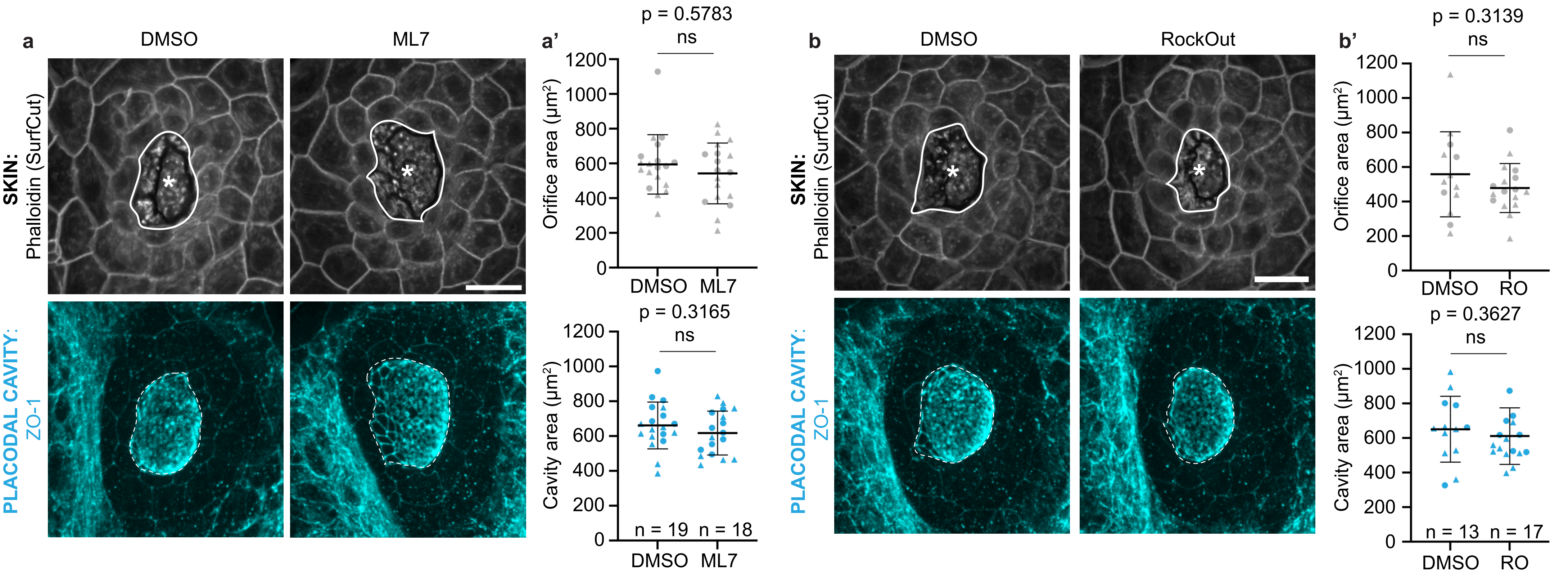

**Supplementary Figure 3. Additional data on the pharmacological inhibition of myosin II phosphorylation,** related to Fig. 4**. a.** Images of 48 hpf embryos treated with ML7 or DMSO from 32 hpf. Top images = skin labelled with phalloidin and extracted with the SurfCut plugin (see Methods). Asterisk = orifice. White line = orifice contour. Bottom images = cavity seen with ZO-1 immunostaining, maximum projections. Dotted white line = cavity contour. Scale bar = 20 µm. **a'**. Quantification of orifice and cavity area in ML7-treated embryos and DMSO-treated controls at 48 hpf (2 independent experiments, n = 18 ML7-treated embryos, n = 19 DMSO-treated controls). Mann Whitney test for orifice area and unpaired two-tailed t-test for cavity area. **b.** Images of 48 hpf embryos treated with Rockout or DMSO from 32 hpf. Top images = skin labelled with phalloidin and extracted with the SurfCut plugin (see Methods). Asterisk = orifice. White line = orifice contour. Bottom images = cavity seen with ZO-1 immunostaining, maximum projections. Dotted white line = cavity contour. Scale bar = 20 µm. **b'**. Quantification of orifice and cavity area in Rockout-treated embryos and DMSO-treated controls at 48 hpf (2 independent experiments, n = 17 Rockout-treated embryos, n = 13 DMSO-treated controls). Welch's t-test for orifice area and Mann Whitney test for cavity area.

Supplementary Figure 4 Gordillo Pi et al.

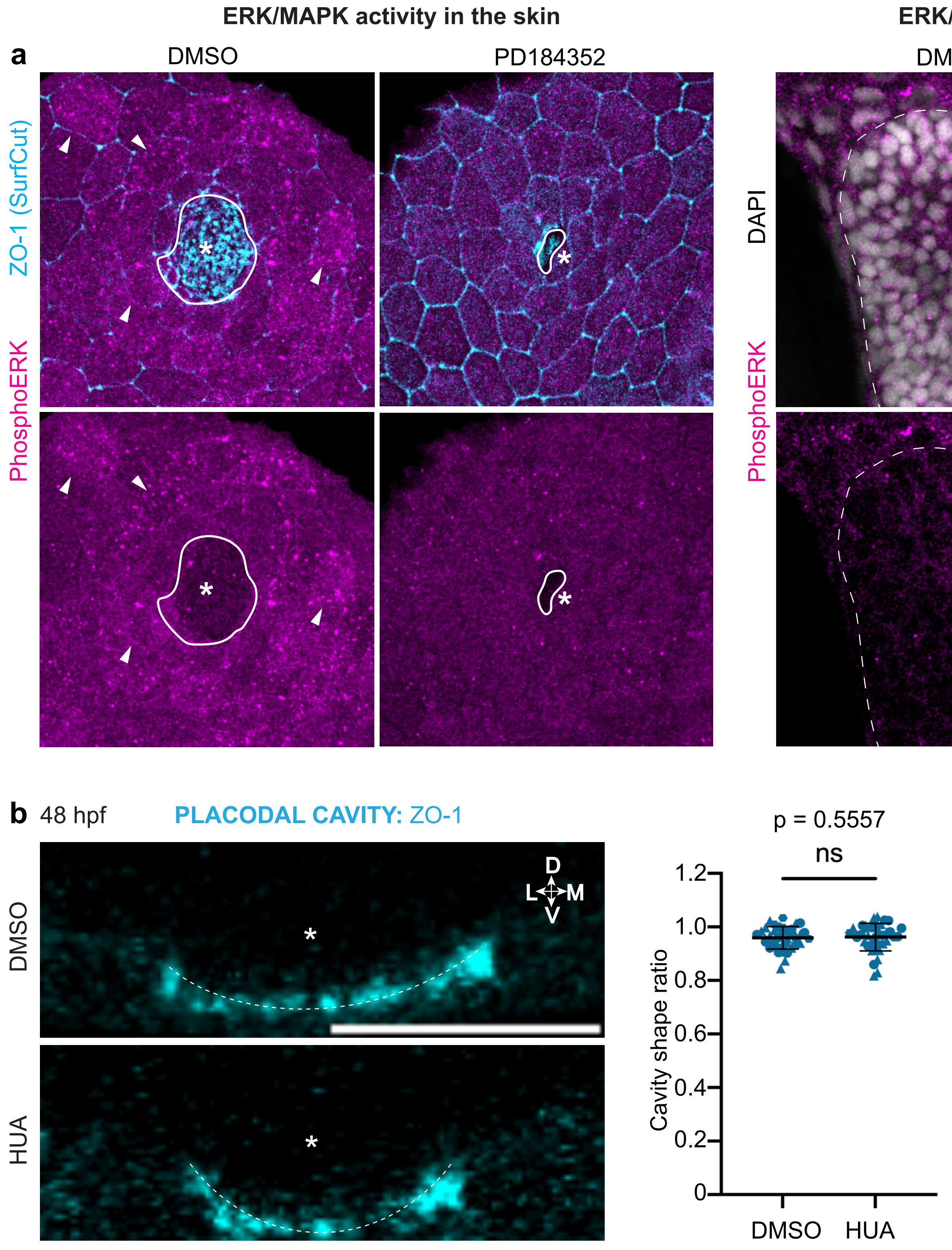

**Supplementary Figure 4. Additional data on the pharmacological inhibition of ERK signaling,** related to Fig. 5**. a.** Left. ZO-1 (cyan) and phospho-ERK (magenta) immunostaining in the skin, extracted with the SurfCut plugin (see Methods), in 48 hpf embryos treated with PD184352 or DMSO from 32 hpf. Arrowheads point to skin cells showing Phospho-ERK immunoreactivity in DMSO-treated controls. This staining is lost in PD184352-treated embryos. Asterisk = orifice. White line = orifice contour. Scale bar: 20 µm. Rigth. Phospho-ERK immunostaining (magenta) and DAPI staining (grey) in the olfactory placode in 48 hpf embryos treated with PD184352 or DMSO from 32 hpf. Dotted line = placode contour. Arrowheads point to olfactory placode cells showing Phospho-ERK immunoreactivity in DMSO-treated controls. This staining is lost in PD184352-treated embryos. Scale bar: 20 µm. **b.** Left. Orthogonal sections (reslice from the z-stack) of DMSO and HUA-treated embryos immunostained for ZO-1 (cyan) to reveal the cavity at 48 hpf. Dotted line = cavity contour. Asterisk = orifice position. Scale bar = 20 µm. Right. Quantification of the cavity shape ratio (as defined in Fig. 5b''') in HUA-treated embryos and DMSO-treated controls (3 independent experiments, n = 39 HUA-treated embryos, n = 37 DMSO controls). Mann-Whitney test.
